# A live cell imaging-based assay for tracking particle uptake by clathrin-mediated endocytosis

**DOI:** 10.1101/2024.02.08.579544

**Authors:** Grant Ashby, Kayla E. Keng, Carl C. Hayden, Jeanne C. Stachowiak

## Abstract

A popular strategy for therapeutic delivery to cells and tissues is to encapsulate therapeutics inside particles that cells internalize via endocytosis. The efficacy of particle uptake by endocytosis is often studied in bulk using flow cytometry and Western blot analysis and confirmed using confocal microscopy. However, these techniques do not reveal the detailed dynamics of particle internalization and how the inherent heterogeneity of many types of particles may impact their endocytic uptake. Toward addressing these gaps, here we present a live-cell imaging-based method that utilizes total internal reflection fluorescence (TIRF) microscopy to track the uptake of a large ensemble of individual particles in parallel, as they interact with the cellular endocytic machinery. To analyze the resulting data, we employ an open-source tracking algorithm in combination with custom data filters. This analysis reveals the dynamic interactions between particles and endocytic structures, which determine the probability of particle uptake. In particular, our approach can be used to examine how variations in the physical properties of particles (size, targeting, rigidity), as well as heterogeneity within the particle population, impact endocytic uptake. These data impact the design of particles toward more selective and efficient delivery of therapeutics to cells.

## ARENA

### Physical Situation

Targeted therapeutic delivery has been an aim in pharmaceutical research for the past three decades (Harrington et al., 2000). Cells employ multiple internalization pathways that can be targeted through modulation of physical particle properties and inclusion of ligands that can target membrane-associated receptors (Eroğlu & İbrahim, 2020). One of the best characterized endocytic pathways, clathrin-mediated endocytosis, is often targeted for therapeutic delivery because it is known to recycle key receptors that are modulated in disease, such as transferrin-receptor (Ohno et al., 1995; Traub, 2009), low-density lipoprotein receptor (Davis et al., 1987; Traub, 2009), receptor-tyrosine kinases (Goh & Sorkin, 2013), and G-protein coupled receptors (Bhagatji et al., 2009; Traub, 2009). Clathrin-mediated endocytosis occurs through the assembly of adaptor proteins that bind to cytosolic receptor motifs (Maxfield & McGraw, 2004) and enable clathrin oligomerization into a growing clathrin-coated vesicle (Robinson, 2015). Trimers of the coat protein, clathrin, assemble into an oligomeric cage around a membrane vesicle until the GTPase scission protein, dynamin (Antonny et al., 2016; Loerke et al., 2009), arrives to cleave the neck of the vesicle, releasing it into the cytosol for transport and uncoating (Kirchhausen et al., 2014).

Live cell imaging-based methods for tracking the uptake of virus particles have been previously established (Cureton et al., 2010, 2012). However, this approach has been largely restricted to tracking a small number of particles manually. In contrast, little has been done to study how drug-carrier particles, such as liposomes, impact the dynamics of endocytosis. Here, we expand on previous work by applying live-cell imaging and tracking algorithms to monitor the uptake of a large and heterogeneous population of particles in parallel by endocytosis. Our data highlight the impact that variation in the physical properties of particles has on endocytosis. In particular, we find that, the population of particles internalized by cells may have distinct properties from the population that initially came into contact with the cell, suggesting that endocytic pathways select for specific physical properties during uptake. These principles can be used to design therapeutic particles that are taken up by cells more efficiently and specifically. Additionally, these data can inform basic biophysical understanding of endocytosis.

### Challenges

While multiple labs have studied endocytic uptake of individual viral particles (Mazel-Sanchez et al., 2023) using TIRF microscopy and lattice-light sheet microscopy (Joseph et al., 2022), these techniques have not been adapted to monitor the uptake of synthetic particles, such as liposomes and beads, commonly used for therapeutic delivery. Tracking the uptake of individual particles is challenging because it is difficult to differentiate between those that are internalized and those that associate with the cell surface but ultimately dissociate from it without becoming internalized. Using TIRF microscopy to examine internalization events at the basal, adherent surface of the cell helps to overcome this challenge. Specifically, the evanescent TIRF field will only illuminate particles within ∼200 nm of the adherent cell surface (Fish, 2009), effectively isolating particles that are diffusing beneath from those in the surrounding solution. Within this environment, particles are effectively confined to a two-dimensional plane, slowing their diffusion, such that they can be reliably tracked. Then, when a particle disappears from the TIRF field simultaneously with an endocytic structure, it can be reasonably assumed that the particle has been internalized by the cell via endocytosis.

To successfully track internalization of particles using this method, the particles must be bright enough to be tracked over a time-scale of at least a few minutes without substantial loss of fluorescence signal owing to photobleaching. Minimizing photobleaching is particularly important for smaller particles that likely contain fewer fluorescent labels. An additional challenge is to prevent particles from adhering to the coverslip to which cells are adhered. Adsorption of particles to this surface limits the pool of particles available to interact with the cell, making it difficult to study the internalization process. Passivating the surfaces of particles with hydrophilic molecules such as polyethylene glycol chains helps to minimize adhesion of particles to the substrate.

### Typical Solution(s)

While the membrane trafficking field has utilized fluorescent microscopy techniques to study endocytosis for decades, these methods have rarely been applied to elucidate the uptake of therapeutic particles (Robinson, 2015). In particular, recent advancements in TIRF microscopy and high-resolution imaging are largely employed to study the basic molecular mechanisms of endocytosis and membrane traffic (Day et al., 2021; Nawara et al., 2022). In contrast, pharmaceutical scientists, who tend to focus on the overall efficiency of particle uptake, more commonly employ batch techniques such as flow cytometry (Sklar et al., 2007) and Western blot (Heath et al., 2016) to judge the efficacy of their therapeutic carriers. Using these techniques, the endocytic pathway responsible for internalization can be characterized by treating cells with small molecule inhibitors of specific pathways, such as clathrin inhibitors (Dejonghe et al., 2019) and dynamin inhibitors (Kirchhausen et al., 2008). If there are decreases in the overall uptake when the clathrin pathway is inhibited as indicated by altered fluorescent signal when analyzed by flow cytometry, or altered protein expression when using Western blots, then it can be assumed that the altered pathway must be a significant route for internalization of the particle. Often these results are confirmed through conventional confocal microscopy, which can assess the extent to which labeled particles appear in cellular endosomes and in the cytosol (Kim et al., 2022).

### Common issues

While these batch methods provide much valuable data, they cannot elucidate the mechanics of particle uptake or the impact of particle heterogeneity. Specifically, because these methods do not track individual particles, they cannot provide information about the dynamics of individual uptake events (Huth et al., 2006). Importantly, most methods for preparing therapeutic particles yield a heterogeneous distribution of particles in terms of variable particle diameter (Gkionis et al., 2020) and chemical composition (Larsen et al., 2011). Determining which particle properties yield the most efficient internalization may provide important insights into the design of more effective particles. However, determining which particles within a heterogeneous batch are taken up most efficiently requires a method that tracks individual particles. Additionally, many of the inhibitors used to reduce uptake through endocytic pathways lack the required specificity to pinpoint an individual pathway. For example, dynamin inhibitors lack specificity owing to the involvement of dynamin in many different routes of endocytosis (Ferguson & De Camilli, 2012). Similarly, drugs such as cyclodextrin, which deplete the plasma membrane of cholesterol (Zidovetzki & Levitan, 2007), are commonly employed to reduce particle uptake by caveolae, yet are known to have broad impacts on cell physiology, which likely also impact uptake through diverse pathways (Rusznyák et al., 2021).

### Method for High Throughput Tracking of Individual Particles During Cellular Uptake via Clathrin-Mediated Endocytosis

#### Big idea

Here we present a method to track the uptake of hundreds of individual particles simultaneously during clathrin-mediated endocytosis. This method combines TIRF time-lapse microscopy, which restricts imaging to the adherent surface of mammalian cells, and quantitative image analysis, which finds and tracks individual particle trajectories within TIRF movies.

Using TIRF microscopy, we perform a 3-channel fluorescence imaging experiment in which we track the dynamics of fluorescently labeled components of the endocytic machinery, which internalize a fluorescently labeled model receptor, along with fluorescently labeled particles, which associate with the receptor at the cell surface (Figure 1A-B). For the purpose of this demonstration, we engineered a model receptor which, as described below, is predominately internalized by the clathrin-mediated endocytic pathway and has a specific affinity for an engineered ligand on the surfaces of the particles we are studying (Figure 1A). However, this method can be adapted to study other particle types, receptors, and endocytic pathways. The basic requirements are orthogonal fluorescent labeling of: (i) particles, and (ii) one or more key protein components of the endocytic pathway under study. Additionally, if targeted delivery of particles is desired, a receptor-ligand system may be employed, as described here, where the ligand resides on the particle surface and the receptor is labeled in a third, orthogonal fluorescence channel.

**Figure 1.**
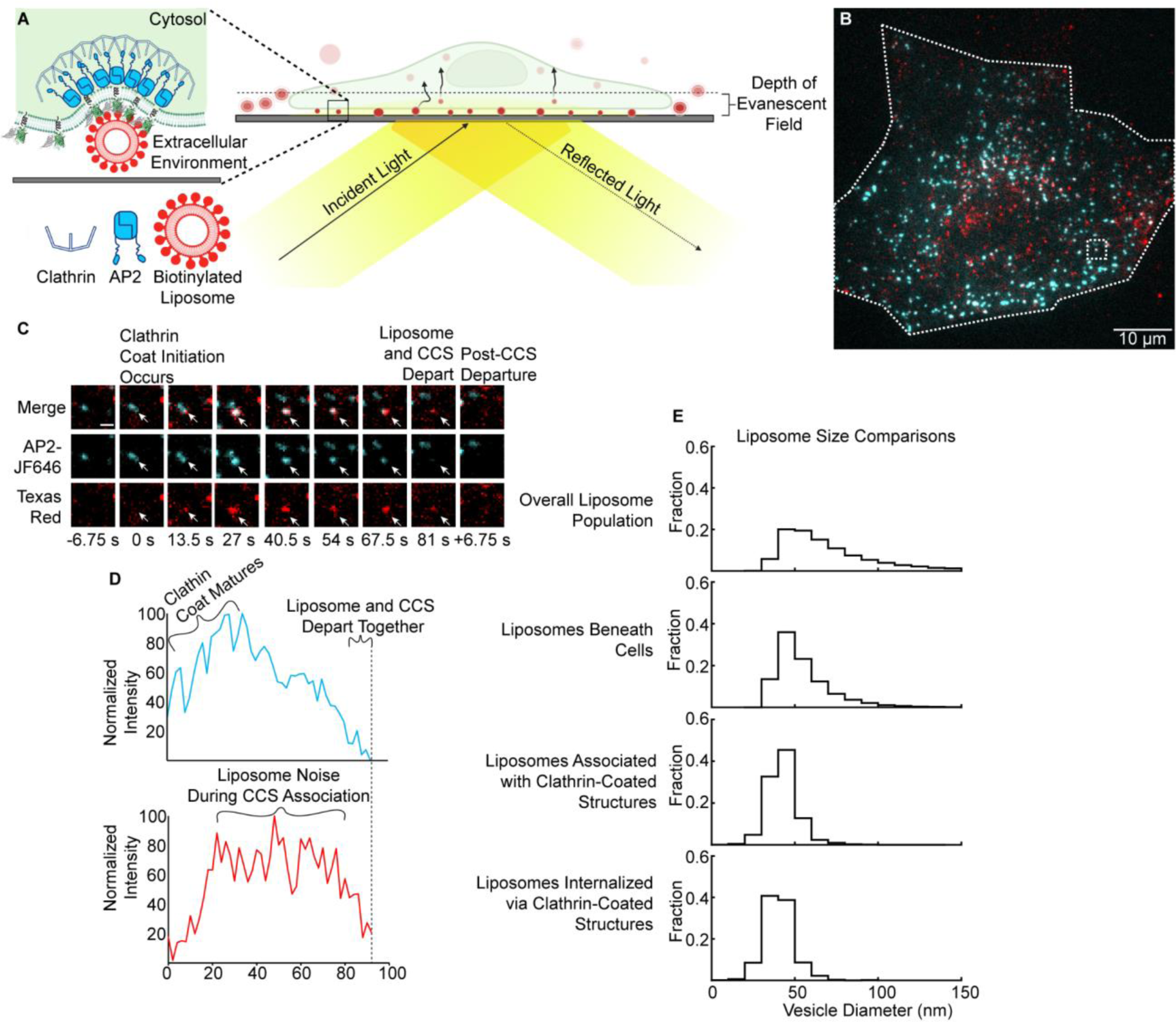
Overall workflow for how individual particles can be judged for internalization and averaged to inform on optimal particle properties for uptake. **A.** A schematic of the basolateral membrane of a cell expressing a target receptor can be analyzed via TIRF microscopy. (Created with BioRender) **B.** An example image of a SUM159-AP2-σ-Halotag cell in the presence of model drug-carriers beneath the basolateral membrane. The endocytic marker, AP2, is shown in cyan, and the model drug-carrier, is shown in red. The periphery of the cell is highlighted by the dashed line, and the dashed inset shows a model colocalization event. **C.** A montage of the growing endocytic site shown in the dashed box in Figure 1B as it interacts with a model drug-carrier through time. The scale bar is 1 µm. **D.** The tracked intensity of the growing endocytic site shown in Figure 1C, as it matures and departs from the cellular membrane with the simultaneous decrease in fluorescent intensity indicating successful drug-carrier internalization. **E.** Normalized histograms of the drug-carrier sizes that were made (47,502 events), penetrated beneath cells (26,537 events), associated with endocytic machinery (15,152 events), and were successfully internalized (1,797 events). Histograms were populated across 3 trials with at least 12 cells analyzed per trial for a total of 84 cells exposed to DOPC liposomes containing 10% biotinylated lipids.

We analyze TIRF movies to identify endocytic sites that internalize particles, which are evidenced by a simultaneous decrease in the fluorescence intensity of all fluorescent channels (Figure 1C-D). Using automated software analysis, we can analyze hundreds of these events per cell, creating relatively large data sets. From these data sets, statistics can be calculated which reveal the impact of variation in the physical properties of particles (diameter, rigidity) on their probability of internalization. For example, Figure 1E compares the distribution of particle diameters for particles that (i) are exposed to cells, (ii) penetrate beneath adherent cells, (iii) associate with endocytic structures, and (iv) become internalized by endocytic structures. These data indicate that smaller particles have a higher probability of overcoming successive barriers to uptake (penetration, association with endocytic machinery) and ultimately being successfully internalized, likely owing to the lesser steric barrier that they pose to the cell’s endocytic machinery. This approach can, in principle, be applied to the wide variety of therapeutic carriers described in the literature, many of which are intended for internalization by cells via endocytic pathways.

## BEFORE YOU BEGIN

### Buffer Preparation

#### Timing: Prior to Starting

The first step is to prepare the particle suspension that will be added to the cells. Depending on the particle type, different buffers will be used, which should be prepared in advance:

One type of particle that we have used to validate our protocol is a population of small liposomes. Liposomes are created by hydrating a dried lipid film with a buffer, as described below. We have had good results using the following buffer for this purpose: 25 mM HEPES, 125 mM NaCl, pH, 7.4. Prepare this buffer if you intend to work with liposomes.

A second type of particle that we have used to validate our protocol is polystyrene beads. To prepare the beads for the experiment, a buffer is needed to dilute the beads and thereby prevent their aggregation. The following buffer works well in this application: 125 mM NaHCO_3_, 20 mM KCl, 1 mM tris(2-carboxyethyl) phosphine (TCEP), pH 8.2. Prepare this buffer if you intend to work with beads.

Both buffers should be vacuum filtered to remove particulates through a 0.2 μm filter.

### Valap Preparation

#### Timing: Prior to Starting

During live cell imaging, samples must be sealed to prevent evaporation. For this purpose, we recommend preparing a batch of “Valap”, a sealant that melts at relatively low temperature (∼80C) and solidifies within seconds, such that it is highly convenient for microscopy applications. Valap stands for **Va**seline: **La**nolin: **Paraffin**. Weight out about 100 g of each into a glass container, heating and stirring until melted and well mixed. Allow to solidify and store at room temperature for repeated use. Re-melt using a hotplate and apply in thin layers with a small paintbrush to seal slides, as described below. Valap will resolidify within seconds upon application in thin layers.

### Preparation of PLL-PEG-Biotin, a reagent used to tether biotinylated particles to coverslip surfaces during particle size calibration

#### Timing: Prior to Starting

A chimera of Poly-lysine, polyethylene glycol, and biotin (PLL-PEG-biotin) is used to tether biotinylated particles, such as liposomes, to the surfaces of coverslips so that the distribution of their fluorescence intensities can be measured. This distribution is then compared to a distribution of particle diameters (measured by dynamic light scattering as described below), to determine a conversion factor between particle fluorescence and diameter. Using this conversion factor, it is possible to estimate the diameters of individual particles during endocytic uptake experiments. The protocol for creating PLL-PEG-biotin has been previously published (Snead & Stachowiak, 2018) and is summarized here.

The purpose of PLL-PEG-biotin is to link particles, via biotin-streptavidin interactions, to the surfaces of ultraclean coverslips. Here PLL, which has a net positive charge, binds non-covalently to the negatively charged glass surface. PEG serves as a neutral, hydrophilic spacer, while biotin provides a specific linkage to streptavidin, which also binds to biotin moieties on the surfaces of the particles.

To form PLL-PEG-biotin, first dissolve poly-L-Lysine (PLL) powder in 50 mM sodium tetraborate pH 8.5 to a final concentration of 50 mg/mL. Calculate the amount of methoxypoly(ethylene glycol) succinimidyl valerate (mPEG-SVA) required for a molar ratio of one mPEG-SVA monomer to every five PLL subunits. To promote binding of biotinylated particles, we include a 1:50 ratio of biotin functionalized PEG that can create a “biotin-sandwich” if neutravidin, which has four biotin binding domains, is introduced. Specifically, biotin-PEG-SVA should be added at a 1:50 ratio to the PEG-SVA. Follow the steps below to create PLL-PEG-biotin:

1. The PEG-powders should be weighed and combined within a nitrogen-filled glove box to prevent hydrolysis.
2. The PLL powder should then be added to the PEG-powders.
3. Dissolve by adding 1 mL of 50 mM sodium tetraborate, pH 8.5 buffer, and vortex to mix.
4. Spin the resulting solution in a test tube with a small stir bar at room temperature for a minimum of six hours.
5. After stirring, size-exclusion columns (Zeba, ThermoFisher, see table below) hydrated with a Hepes buffer (25 mM HEPES buffer, 125 mM NaCl, pH 7.4) should be used to remove unreacted poly-l-lysine.

### Cell Line Selection

#### Timing: Prior to Starting

The cell line chosen for this assay should express a bright, highly specific marker for the endocytic pathway of interest, which, in our case, is clathrin-mediated endocytosis. We have chosen to use SUM159 cells, a breast tumor-derived epithelial cell line, which has been gene-edited at the endogenous locus in both alleles of the sigma2 domain of adaptor protein 2 (AP2) to include a HaloTag domain at its C-terminus (SUM159-AP2-σ2-Halotag) (Aguet et al., 2016). These cells were generously provided by the T. Kirchhausen laboratory (Harvard University). AP2 is the major adaptor protein of the clathrin pathway(McMahon & Boucrot, 2011). It binds simultaneously to the cytosolic motifs of transmembrane proteins, PI(4,5)P_2_ lipids, and clathrin triskelia, helping to create clathrin-coated vesicles that are loaded with transmembrane cargo (Collins et al., 2002; Kelly et al., 2014). AP2 is found in approximately 1:1 stoichiometric ratio with clathrin (Cocucci et al., 2012), making it a good marker for tracking the growth and dynamics of clathrin mediated endocytic structures. SUM159 cells are also optimal for imaging owing to their wide, thin lamellipodia (Aguet et al., 2016), which help to minimize background signals during TIRF microscopy.

Once the cell line and particle have been selected, we recommend trial imaging of particle delivery to cells over 10 to 15 minutes to access: i) lack of spectral overlap between the chosen fluorescent markers, ii) photo-stability of the labeled particle and endocytic components, and iii) sufficient penetration of particles beneath cells. If these criteria are met, then TIRF imaging can proceed.

### Cell Culture Media Preparation

#### Timing: Prior to Starting

For culture of the SUM159-AP2-σ2-Halotag cells, the following buffers should be prepared:

Buffered Maintenance Media: 1:1 mixture of DMEM high glucose and Ham’s F-12 media, supplemented with 5% v/v fetal bovine serum (FBS), 10 mM HEPES, 5 μg/mL insulin, 1 μg/mL hydrocortisone, and 1% v/v penicillin/streptomycin/L-glutamate, pH 7.4.

<u>Transfection Media without FBS</u>: Buffered Maintenance Media lacking phenol-red pH indicator, FBS, and antibiotic. FUGENE HD was used for transfection of cells with the model receptor.

<u>Transfection Media with FBS:</u> Cells require serums for optimal growth and proliferation. For this reason, we need to create a transfection media mixture that also contains FBS. This mixture should contain a 1:1 mixture of DMEM high glucose and Ham’s F-12 Media but should omit the phenol-red pH indicator. This mixture should be supplemented with 5% v/v FBS, 10mM HEPES, 5 μg/mL insulin, 1 μg/mL hydrocortisone, pH7.4. Notably, this media lacks antibiotics, which reduce transfection efficiency.

### Acid Washed Coverslip Preparation

#### Timing: Prior to Starting

For plating cells onto coverslips for TIRF imaging, follow the steps below:

1. Separate, and place 30 to 40 coverslips (22 mm x 22 mm 1.5 mm thick) into a 250 mL glass beaker.
2. Add 200 mL of 1M HCl and heat the beaker to 40-60℃ overnight in a fume hood.
3. Cool the beaker to room temperature and rinse out the 1M HCl with ultrapure water.
4. Fill the beaker with 200 mL of ultrapure water and sonicate in a bath sonicator for 15 minutes. Repeat this step two additional times.
5. Fill the beaker with 200 mL of 50% ethanol/50% ultrapure water and sonicate in a bath sonicator for 15 minutes.
6. Fill the beaker with 200 mL of 70% ethanol/30% ultrapure water and sonicate in a bath sonicator for 15 minutes.
7. Fill the beaker with 200 mL of 95% ethanol/5% ultrapure water and sonicate in a bath sonicator for 15 minutes.
8. Fill the beaker with 200 mL of 95% ethanol/5% ultrapure water, cover with aluminum foil, and place in a BSL-2 laminar flow hood.

## KEY RESOURCES TABLE

| REAGENT or RESOURCE | SOURCE | IDENTIFIER |
| --- | --- | --- |
| Chemicals, and Lipids for Liposomes |  |  |
| HEPES Powder | Fisher BioReagents | BP310-1 |
| NaCl | Fisher Chemical | S271-3 |
| Sodium Bicarbonate | Sigma-Aldrich | S6014-500G |
| KCl | Sigma-Aldrich | P5405-1KG |
| TCEP | Sigma-Aldrich | 646547 |
| DOPC | Avanti Polar Lipids | 850475P-200mg |
| DSPE-PEG2K | Avanti Polar Lipids | 880120P-200mg |
| PE-CAP-Biotin | Avanti Polar Lipids | 870273P-25mg |
| Texas Red-DHPE | AAT Bioquest | 23300 |
| DPPC | Avanti Polar Lipids | 850355P-200mg |
| Cholesterol | Avanti Polar Lipids | 700000P-500mg |
| Glass Test-Tubes for Lipid Drying | KIMBLE | 73500-1275 |
| Forced Air Ovens | VWR | 89511-410 |
| RotoVap or Vacuum Pump for drying | KNF Group | LABOPORT- UN811KVP |
| Yellow-Green Biotin FluoSpheres (40 nm) | Invitrogen | F8766 |
| UV cuvette micro | Brand | BR759200 |
| Hamilton "Gastight" Syringes- 10 μL | Hamilton | 80000 |
| Hamilton "Gastight" Syringes- 100 μL | Hamilton | 81000 |
| **Reagents for Tethering Vesicles** |  |  |
| 1.5H glass coverslips (24 x 50 mm) | Thor Labs | CG15KH |
| Poly-L-Lysine | Sigma-Aldrich | P2658-200mg |
| Biotin-PEG-SVA | Laysan Bio | Biotin-PEG-SVA-5000-100mg |
| MPEG-SVA | Laysan Bio | mPEG-SVA-5000-1g |
| Hellmanex III | Sigma-Aldrich | Z805939 |
| Zeba Spin Desalting Columns, 7K MWCO, 5 mL | ThermoFisher | 89891 |
| Neutravidin | ThermoFisher | 31000 |
| 1.6 mm Clear Silicone Sheet | Grace Bio-Labs | 664273-S |
| **Reagents for Cell Culture Media** |  |  |
| Insulin | Sigma-Aldrich | I6634-50MG |
| Hydrocortisone | Sigma-Aldrich | H4001-1G |
| Fetal Bovine Serum | Sigma-Aldrich | F0926-500ML |
| Penicillin/Streptomycin/L-glutamine | Cytiva | SV30082.01 |
| Ham's F-12 | Cytiva | SH30026.FS |
| DMEM High Glucose | Cytiva | SH30285.FS |
| DMEM/F-12 1:1 with L-glutamine, without HEPES and phenol-red | Cytiva | SH30272.02 |
| OxyFluor | Oxyrase | OF-0005 |
| 100 mm Tissue Culture Dishes | Corning | 8-772-23 |
| Costar 6-Well Tissue Culture Treated Well Plate | Fisher Scientific | 07-200-83 |
| **Critical Commercial Assays** |  |  |
| Zetasizer NanoZS | Malvern Instruments | MAN0485-1-1 |
| NanoDrop ONE-C | ThermoScientific | ND-ONE-W |
| SLPe Digital Sonifier | Branson | SLPe 40:0.15:4C |
| **Experimental Models: Cell Lines** |  |  |
| SUM159-AP2-σ2- HaloTag | Aguet (lattice) & Kirchhasen | N/A |
| **Recombinant DNA** |  |  |
| TfR-mEGFP-mSA | Ashby & Stachowiak | N/A |
| TfR-mRFP-mSA | Ashby & Stachowiak | N/A |
| **Software and Algorithms** |  |  |
| MATLAB 2017B | Mathworks | N/A |
| CMEanalysis | Danuser & Aguet | N/A |
| Internalization Filters | Ashby & Stachowiak | N/A |
| **Reagents for Cell Coverslip Imaging** |  |  |
| Double-Sided Tape | Scotch | CBGNHW011091 |
| Acid-Washed Square Glass Coverslips (22 x 22 mm) | Thor Labs | CG15CH |
| Tweezers | VWR | 89411-752 |
| Single-Blade Razor | VWR | 55411-055 |
| Valap Sealant(1:1:1, vaseline: lanolin: paraffin) | Stachowiak | N/A |

## EQUIPMENT

In order to follow the below procedures to image fluorescent particle delivery the following equipment is required:

Equipment for liposome preparation:

● Solvent resistant syringes for mixing lipids dissolved in organic solvents.
● Vacuum chamber for drying of the lipid film.
● A method for multilamellar vesicle disruption into small unilamellar vesicles, some options include:
`o` Probe sonication
`o` Membrane extrusion
● Table-top microcentrifuge for removal of aggregates from samples after sonication or extrusion.
● Ultraviolet/visible spectroscopy system for quantification of the final concentration of probe lipids within the liposome samples (a proxy for total lipid concentration).
● A dynamic light scattering system for quantifying the distribution of particle diameters.

Equipment for preparing biotinylated beads:

● Bath sonicator for breaking up clustered beads.
`o` Notably, the manufacturer states that probe sonication can damage these beads, so bath sonication is the recommended homogenization method.

Equipment for cell-culture:

● BSL-2 laminar flow hood for sterile cell plating and passaging.
● Sterile CO_2_ incubator for mammalian cell growth.

Equipment for cellular-imaging:

● Inverted fluorescence microscope with TIRF illumination.
● A minimum of 3 fluorescent channels.
● Cell-heating capability during imaging.
● Auto-focus capability to maintain focal plane during imaging.

Software for image analysis:

● Particle Tracking Algorithm

We use openly available CMEanalysis which integrates with MATLAB (Mathworks). CMEanalysis is an openly available software package from the Danuser laboratory (Aguet et al., 2013). It is available at the following link: https://github.com/DanuserLab/cmeAnalysis

### Alternative Imaging Methods

Alternative microscopy schemes may be able to achieve similar results. Whereas TIRF microscopy uses an evanescent field that decays within roughly 200 nm of the focal plane (Fish, 2009), spinning-disc confocal microscopy could be applied as well. The axial resolution is not as narrow as TIRF microscopy (roughly 500 nm) (Fouquet et al., 2015), so internalized particles that are not immediately transported away into the cytosol, may be difficult to distinguish from those that are associated with an endocytic structure. Lattice-Light sheet microscopy and structured illumination microscopy likely exceed the necessary temporal and spatial resolution for tracking vesicle uptake (Chen et al., 2014) and may yield interesting results obscured by the diffraction limit in our studies. However, as CMEanalysis was designed around TIRF and confocal image data, data analysis methods may require modification if other imaging methods are used.

## STEP-BY-STEP METHOD DETAILS

### Vesicle Preparation and Characterization

#### One Day Prior to Imaging

1. Rinse round-bottom test tubes (12 mm diameter, 75 mm length) with 3 washes of acetone, 3 washes of water containing 2% detergent (Neutrad), and 3 washes of ultrapure water. Then allow tubes to dry for at least 30 minutes in an oven at 100°C.
2. Remove aliquots of lipid solutions dissolved in chloroform from −80°C freezer and allow to equilibrate to room temperature (25°C) before opening.
3. Mix lipids dissolved in chloroform at desired mole ratios in the round-bottom test tube.
• For liposomes of low membrane rigidity our mole ratios were as follows: 78 mol% DOPC, 10 mol% DOPE-CAP-Biotin, 2 mol% DSPE-PEG2K, and 10 mol% Texas Red DHPE
• For liposomes of high membrane rigidity, our mole ratios were as follows: 68 mol% DPPC, 10 mol% DOPE-CAP-Biotin, 10 mol% Cholesterol, 2 mol% DSPE-PEG2K, and 10 mol% Texas Red DHPE
6. 2. Dry lipids into a thin-film along the bottom of the test-tube with N_2_ gas.
7. Cover the sample with aluminum foil and place inside a vacuum chamber for a minimum of 3 hours.

#### Day of Imaging

1. Resuspend the thin lipid film with 2 mL of HEPES buffer added to each tube. Vortex vigorously to remove the lipid from the walls of the test tube. Then cover and place on ice to further hydrate the lipid films for at least 30 minutes.
2. Vortex again prior to sonication, and then sonicate with a probe sonicator (⅛” probe diameter, SLPe SFX150 Branson Ultrasonics) on ice for 1 minute on and 1 minute off. These 1-minute cycles should repeat 6x each for a total time of 12 minutes.

. We recommend using 50-60% sonication amplitude. As probe use increases, sonication amplitude may need to be increased to create similar liposome diameter distributions. We verify this by judging the power output is between 90 and 100 Watts. Settings will likely vary depending on the sonication instrument. In general, the sonication power and amplitude are correct when the liquid in the tube vibrates, but does not bubble, during sonication, and the outside of the tube becomes warm to the touch during each one-minute cycle of sonication, but not too warm to comfortably hold. Some optimization may be required. If DLS analysis (described below) indicates that the liposome diameter remains above 100 nm, repeat sonication with higher power and/or amplitude.

1. 2. After 1 probe sonication run, replace the ice used and repeat the sonication steps in step 2 for an additional 12-minute cycle to bring the total to 24 total minutes.
2. After the 2 sonication runs, fill 2, 1.5 mL microcentrifuge tubes with 900 μL of lipid resuspension, and spin in a tabletop centrifuge set to 4°C at maximum speed for at least 20 minutes. This spin should pellet metal particulates from the sonicator probe and any potentially large lipid aggregates.
3. Remove 750 μL of the supernatant from each microcentrifuge tube and move to a new tube.
4. Take at least 200 μL of the combined tube and place it in a disposable cuvette. Use this cuvette to run Dynamic Light Scattering to determine the diameter distribution of the vesicles after sonication.

### Tethered Vesicle Assay to Convert Particle Fluorescence Intensity to Diameter

To perform the conversion of particle intensity to diameter, first we measure the diameter distribution using dynamic light scattering. Then we measure the distribution of fluorescent intensities by imaging tethered vesicles. These two distributions are then compared to create a conversion factor between fluorescent intensity and particle diameter. This assay leverages the previously published assays from Snead and Stachowiak, and Jensen et. al (Jensen et al., 2011; Snead & Stachowiak, 2018).

#### Dynamic Light Scattering Measurement

1. Use a dynamic light scattering system, such as Zetasizer Nano ZS (Malvern instruments), to acquire the distribution of particle diameters.
2. Load the vesicles into a disposable cuvette with enough sample to fill the cuvette above the minimum volume required for measurement.
3. The Zetasizer NanoZS averages multiple traces together after subsequent runs to collect one distribution of particle sizes. When a proper particle density is present within the cuvette, it is recommended that a minimum of 10 traces are acquired per distribution and for a given group of vesicles that at least 3 distributions are acquired and averaged to get a representative diameter size distribution.
4. The correlation graph should confirm an acceptable polydispersity index with minimal aggregation. Typically, liposome formulations with a polydispersity index below 0.3 are acceptable, meaning that there is a good homogenous size distribution(Danaei et al., 2018). If the measured polydispersity is higher than this value, we recommend repeating the sonication steps listed above.
5. The resulting average, intensity weighted distribution will be used for calibration in subsequent steps.

#### Tethered Vesicle Assay

1. Using tweezers, center a silicone gasket (1.8 mm thick, 4 mm well diameter, Grace Bio-labs) with a minimum of 30 μL well volume onto an ultraclean coverslip. (Figure 2A)
2. Add 12 μL of PLL-PEG-Biotin solution for a minimum of 20 minutes at room temperature to the well within the silicone gasket (Figure 2B).
3. Using the same buffer (HEPES) that you used to prepare the vesicles, wash the coverslip to remove any unbound PLL-PEG-Biotin by adding 60 μL of HEPES buffer a minimum of 8 times. These washes should be performed by gently adding the buffer into the well created by the gasket, then removing that same volume of buffer. Be careful not to touch the glass, as doing so will disturb the layer of PLL-PEG-Biotin that has adhered to the coverslip.
4. Add 4 μL of 1mg/mL neutravidin (resuspended in ultrapure water) to the well. Mix thoroughly by pipetting and allow to incubate for 15 minutes. Neutravidin will bind to the coverslip coated with 2% PLL-PEG-Biotin. Because neutravidin can bind up to 4 biotin moieties, biotinylated particles can be added to bind the additional domains creating a “neutravidin sandwich”.
5. Using the same buffer that you used to make the vesicles, wash the coverslip to remove any unbound molecules from the PLL-PEG-Biotin and neutravidin solution by adding 60 μL of HEPES buffer a minimum of 8 times. Be careful not to touch the glass to disturb the adhered PLL-PEG-Biotin-neutravidin solution.
6. Add 16 μL of diluted vesicle-containing solution to the well. Typical dilution is 1000-5000x.

a. Note that the concentration of purified vesicles might be too high for proper tethering density. On average, vesicles need to be 1-2 diffraction-limited distances apart on the coverslip so that they can be resolved during imaging. If tethering is too dense, the 2D-gaussians will not fit all particles properly, and small particles will be omitted leading to a skewed conversion factor towards large particle sizes.
b. To combat the potential dense particle tethering, it is recommended that a 1000x to 5000x dilution is needed prior to tethering. After dilution, add the vesicle containing solution to each well. Mix thoroughly by pipetting gently up and down upon addition and allow to incubate for a minimum of 15 minutes. Do not introduce air bubbles during pipetting.
7. Using HEPES buffer, wash the well by adding 60 μL of buffer a minimum of 8 times following the same pipette in, pipette out method presented in Step 3. Be careful not to touch the glass, which will disturb the adhered vesicles. Once all the above steps are complete, the sample is ready for imaging.

### Vesicle Imaging to Determine Diameter Distribution

1. Prior to imaging, switch to a widefield setting of the TIRF microscope and ensure all channels in use are focused to a single-point on the ceiling. This test ensures that all channels are entering the microscope body along the same path, as required for TIRF.
2. The microscope should have an autofocus feature to ensure a flat, stable optical plane throughout the experiment.
3. Acquire time-lapse movies of the tethered vesicle well using TIRF microscopy where a minimum of 10 frames are acquired per field of view (Figure 2C). The 10-frame minimum is suggested to allow the detection algorithm to omit any unbound vesicles that are found in one frame but disappear in the next.
4. Acquire a minimum of 1,000 vesicles for proper diameter conversion. On our TIRF microscope, this typically requires a minimum of 10 fields of view to achieve this metric (Figure 2D).

### Vesicle Detection

After the fields of view are acquired, software should be used to identify tethered vesicles, which appear as fluorescent puncta in the fluorescent lipid channel. The software should fit a two-dimensional gaussian function to each diffraction limited-punctum in order to estimate its intensity. Our lab uses an open source software package from the lab of Gaudenz Danuser (UT Southwestern), CMEanalysis, to accomplish this goal (Aguet et al., 2013). We use CMEanalysis in two ways: (i) to detect diffraction limited puncta of low signal-to-noise, and (ii) to track these puncta over time. To analyze tethered vesicles, we will only leverage the detection functionality. Later, in the *Tracking Endocytic Dynamics During Particle Uptake* and *Analyzing Particle Trajectories to Differentiate Among Possible Outcomes* sections, we will leverage both the detection and tracking functionalities to analyze the dynamics of clathrin-mediated endocytosis.

**Figure 2.**
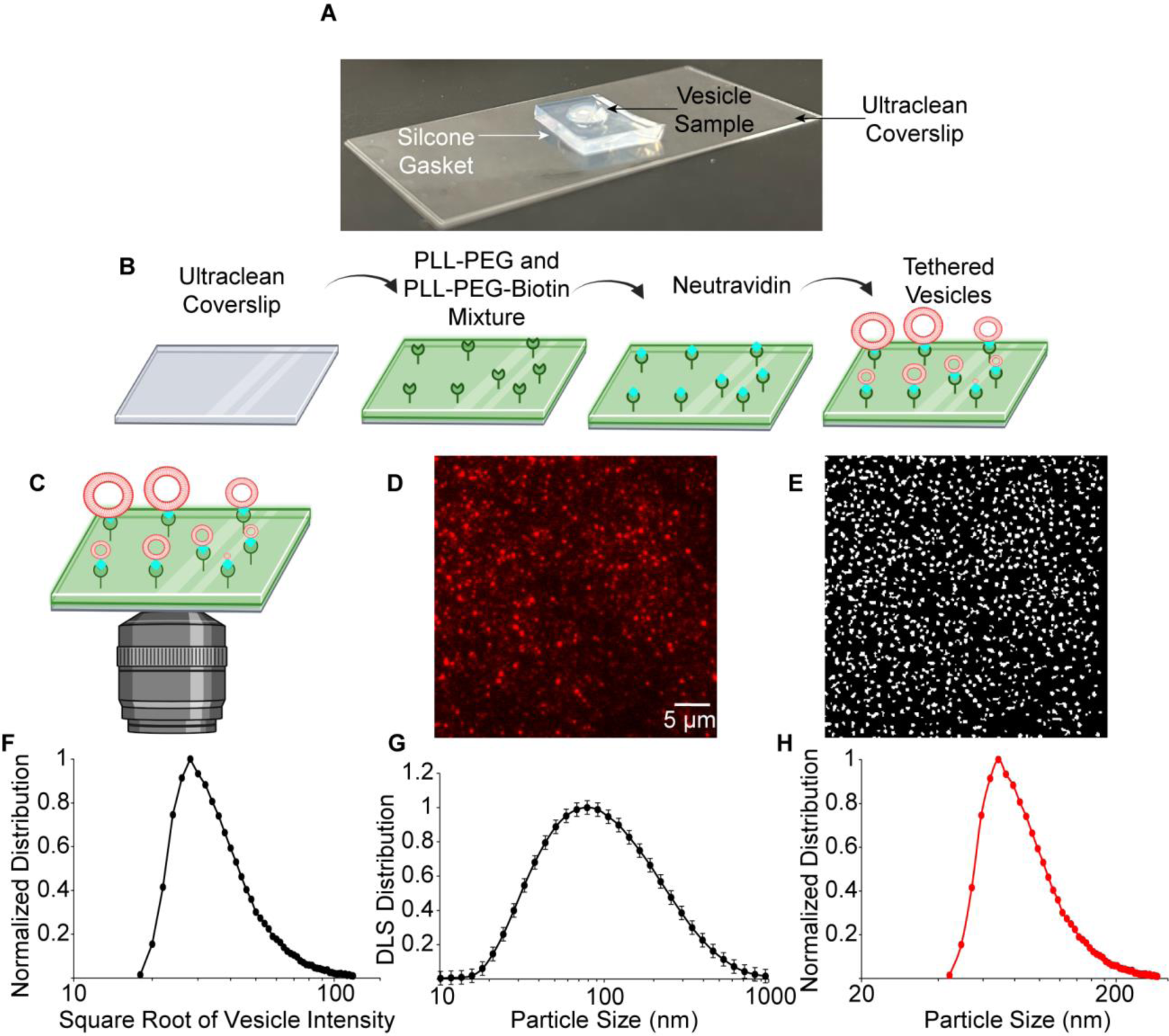
Tethered vesicle assay to enable the conversion of particle fluorescence intensity to diameter. **A)** Snapshot of the experimental set-up of an imaging well on an ultraclean coverslip that is housed by a cleaned silicone gasket. **B)** Schematic of the experimental set-up of how to apply reagents to an ultraclean coverslip to allow for tethering of biotinylated liposomes. (Created with BioRender) **C)** Schematic of the final imaging condition to be acquired under the same imaging parameters that were utilized in cellular delivery. (Created with BioRender) **D)** Representative raw image of tethered DOPC tethered vesicles. **E)** Representative mask of the image in part D from the automated analysis protocol, CME Analysis, to provide the intensity distribution of vesicles present. **F)** Histogram counts of the square root of vesicle intensities from data like D and E. **G)** Representative dynamic light-scattering distribution to provide the range of vesicle diameter created during probe sonication averaged over 3 independent trials. **H)** Converted vesicle diameter utilizing a conversion factor to relate the vesicle intensities to vesicle diameters shown in F and G.

To detect the tethered vesicles, follow these steps:

1. Pass the acquired TIRF images of tethered vesicles through the detection algorithm of CMEanalysis to detect all diffraction-limited puncta in the field of view. Only objects that appear in 3 subsequent frames should be included in the analysis.

a. The amplitude of the two-dimensional gaussian should be stored for fluorescent intensity conversion (Figure 2E). In CMEanalysis the amplitude of the gaussian fit is referred to as an “A-value”, a terminology we will adopt from this point forward in the protocol.
b. CMEanalysis will only store A values that are two standard deviations above the local background. If this criterion is met, a two-dimensional gaussian is fit, and the average intensity of the local background is also stored as a “C-value”.
2. The vector of all A values represents the distribution of fluorescence intensities for the vesicles. The next section describes how to compare this distribution with the distribution of particle diameters (from DLS) in order to create a conversion factor (Figure 2F).

### Vesicle Intensity to Diameter Conversion

1. Compute an intensity scaling factor which will be denoted as f to convert the A values to diameter values where: f=D/√A_median_. D should be the average diameter from the intensity-weighted diameter distribution, and A_median_ should be the median intensity of the A values computed above. Here we assume that the number of labeled lipids per vesicle should scale with the surface area of the vesicle, such that the square root of the vesicle intensity is proportional to the vesicle diameter. The value of D in the above equation corresponds to the peak of the vesicle diameter distribution from DLS (Figure 2G).
2. Using this conversion factor, f, all A values can be multiplied by f to convert diffraction limited particle intensities to particle diameters where: F*√A_particle_ = Diameter_particle_ (Figure 2H).

### Cell Culture and Vesicle Introduction

#### Two Days Prior to Imaging

1. Prepare the required number of acid-washed coverslips.

a. Remove coverslips from the 250 mL beaker containing 95% ethanol.
b. Use heat, or place vertically in a culture dish, to dry the coverslips. 22 x 22 mm coverslips can stand vertically in a 12 or 24 well plate.
2. Prepare cells following standard protocols for cellular detachment and plating in a new culture dish.
3. Resuspend and count the cells.
4. Add the acid-washed coverslips to a 6-well dish containing phenol-red media and add the calculated resuspension volume to add 50,000 cells per well to each well.
5. Return cells to the incubator overnight.

#### One Day Prior to Imaging

1. Aspirate media from the wells and add 900 μL of phenol-red free media containing Fetal Bovine Serum to each well.

The following steps assume FUGENE HD transfection agent is being used. If other transfection agents are being used, the steps may require modification:

1. 2. The transfection cocktail will be composed of FBS free transfection media, FUGENE HD, and plasmid to a final volume of 100 μL per well, such that the total volume in each well is 1 mL. Perform the following for each well to be transfected:
2. Add to a 1.5 mL microcentrifuge tube in the following order:
• Transfection Media without FBS
• DNA plasmid
• FUGENE HD
6. b. For each tube: Add a volume of plasmid such that the final concentration of DNA in the tube is 1000 ng/μL. Add 3μL of FUGENE HD transfection reagent. Calculate the final volume of FBS free transfection media needed to reach 100 μL per well. The volume of media added should be calculated from: 100μL - 3μL (for the volume of FUGENE HD) - the volume of plasmid added. The final volume of media added should typically be between 94 and 90 μL per well.
7. Invert the microcentrifuge tube at least 10 times to mix, tap the tube on a hard surface to send fluid back down to the bottom of the tube, and allow the tube to rest for 15 minutes.
8. After 15 minutes, add 100 μL of the DNA/FUGENE/Media mixture to each well one drop at a time. Then return the 6-well plate to the incubator overnight.

#### Day of Imaging

1. Warm PBS, Transfection Media with FBS, and Oxyrase to physiological temperature.
2. Wash each well with 5 washes of 1 mL of PBS to remove residual FUGENE and replace with 1 mL of Transfection Media with FBS.
3. Use UV/VIS spectroscopy to measure the Texas Red absorption of the liposome solution and calculate the amount of Texas Red in the liposome sample. Multiply this value by 10, assuming 10 mol% Texas Red DHPE was used in the liposomes, to estimate the total concentration of lipids in the sample, as some lipids may have been lost during the vesicle formulation process.
4. Using the total lipid concentration from the concentrated stock, calculate the volume of lipid required to create a final 1 mL well volume containing 1 μM of total lipid.
5. Remove from each well a volume of media equivalent to the volume of lipid solution calculated above and an additional 1.25 μL (to account for the addition of the JF646 Haloligand, which should be dissolved in a solution of DMSO at concentration of 100 μM). Then add the calculated volume of small unilamellar vesicles from the concentrated stock as well as 1.25 μL of the JF646 JaneliaFluor ligand stock to each well.
6. Return the 6-well plate to the incubator for 15 minutes.
7. During this 15-minute incubation, make an imaging media aliquot. This aliquot should contain 1 μM of total lipid and a 1:33 volume/volume ratio of Oxyrase which will aid in removal of oxidative species during fluorescent imaging. The remaining volume should be FBS containing transfection media, according to the recipe above. A minimum volume of 37 μL per coverslip is required.
8. Create a chamber well with fresh media for imaging. To do so, we clean a standard glass microscope slide (25.4 mm x 76.2 mm) with ethanol and apply 2 strips of double-sided tape (∼5mm in length x 1 mm) parallel to the long-axis of the slide, as shown in Figure 3. It is imperative that the double-sided tape is flat against the slide and does not have bubbles (see Figure 3). This is a critical step; errors in this step are the most common reason for failed TIRF imaging. After tape application, use a razor blade to remove excess over the edge of the slide, and move the tape-covered slide to the cell culture hood. A list of further potential sample preparation failures and troubleshooting suggestions are listed in the *OPTIMIZATION AND TROUBLESHOOTING* section below.
9. After 15 minutes, remove the 6-well plate from the incubator and aspirate the media from the desired well. Using tweezers, stand the coverslip on its end and hold it along the edge with your gloved thumb and forefinger to allow for drying of the coverslip edge. Drying the edge will allow for tight adherence of the coverslip to the double-sided tape to the mounting slide. Using a clean chem wipe, wipe all 4-top facing edges (side on which cells are plated) and the entire bottom face (side on which cells are not plated). Align the coverslip over the mounting slide and lay the dried edge down (adherent cell side down) against the double-sided tape and press on the glass coverslip against the tape with tweezers to ensure a tight seal (see Figure 3).
10. In the channel well created in the previous steps, add at least 37 μL of oxyrase/liposome/media mixture. Then, using valap heated such that it is melted, seal all edges of the coverslip, slide interface to trap the liquid mixture and prevent evaporation during sample imaging. Avoid creating any air bubbles (Figure 3). Notably, the vesicles added in step 5 above were largely lost during creation of the sample slide. It is for this reason that additional vesicles are added in this step. These extra vesicles will provide a source of material for continuous uptake throughout imaging.

### Cell and Vesicle Imaging

Similar to the two steps listed in “Vesicle Imaging Parameters”: Prior to imaging, switch to a widefield setting of the microscope and ensure all of the laser lines in use are focused to a single-point on the ceiling. This ensures that all channels are entering the microscope along the same parallel path, as required for TIRF. The microscope should have an autofocus feature to ensure a flat, stable optical plane throughout the experiment.

**Figure 3.**
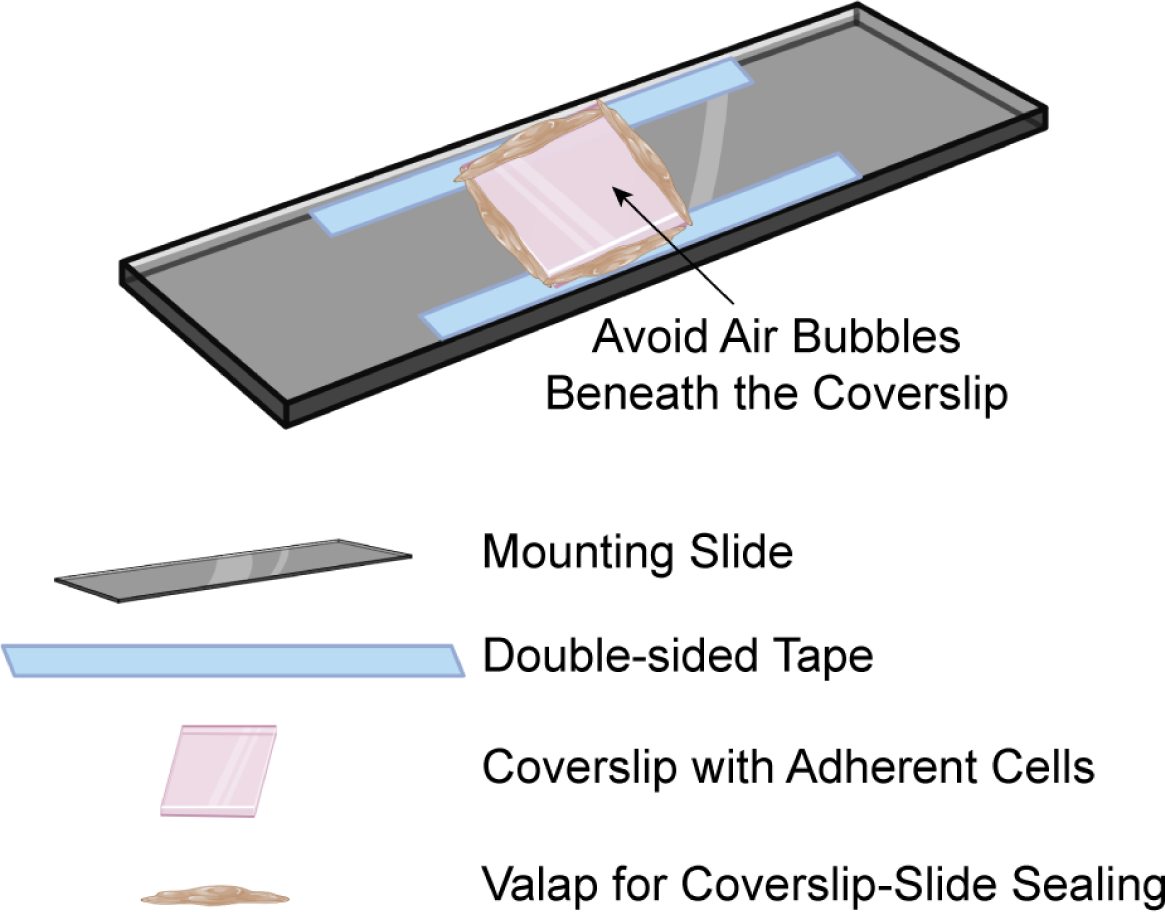
Schematic of cell-mounting slide formation with double-sided tape, coverslip, and valap for sealing. (Created with BioRender).

Beyond the imaging parameters required to image tethered vesicles, the following additional parameters should be noted:

1. The sample should be maintained at 37°C to ensure proper cellular function during the experiment.
2. A frame rate of at least ten times the estimated rate of movement of the particles should be used. It has been previously reported that a 2s to 3s frame rate is sufficient to study endocytic events. To study liposome diffusion we found that a frame rate between 1s and 2s is optimal, based on the mobility of particles diffusing between the cell and the coverslip.
3. With the increased frame rate, there is a risk of increased photobleaching, particularly for smaller particles. Therefore, particles must be labeled with a substantial number of bright, stable fluorophores to ensure visualization throughout the experiment. We found that a minimum of 1000 Texas Red labeled lipids per vesicle were required to sufficiently track vesicles with a diameter of approximately 30 nm (10 mol% of total lipid), the smallest in our experiments.
4. Productive endocytic events occur within 20 to 180 seconds (Loerke et al., 2009). IN our experience, movies of 10-15 minutes in length are sufficient to avoid biasing the analysis toward short-live endocytic events by cutting off a significant fraction of longer trajectories.
5. The signal to noise ratio is relatively low when tracking individual diffraction-limited endocytic structures. Therefore, to reach statistically significant conclusions, thousands of endocytic events must be analyzed. We were typically able to identify a few hundred endocytic events per movie, such that movies of 10 or more cells were required to accumulate a sufficient amount of data per trial. Additionally, we found that each cell sample could only be imaged for about an hour before the cells began to show signs of declining viability, which can impact endocytic dynamics.

### Tracking Endocytic Dynamics During Particle Uptake

After acquisition of live-cell TIRF movies, we employ CMEanalysis to detect the diffraction limited puncta present in every frame of the movies. Here, we employ both the detection and tracking functions of CMEanalysis to examine the dynamics of endocytic events:

1. Pass all acquired TIRF movies through the analysis algorithm to detect diffraction limited puncta in all channels and to track all diffraction limited puncta in the endocytic marker channel (AP2-JF646). In CMEanalysis, only one channel is tracked through time, typically the endocytic marker channel (in our case, AP2-JF646). This tracked channel is denoted as the “master” channel. All other channels are designated as “subordinate channels” and analyzed for overlap with the master channel. In our analysis, the receptor and particle were typically subordinate channels.
2. In the master channel, CMEanalysis fits a 2D gaussian function to each diffraction-limited object that has an intensity that is at least 2 standard deviations above the local background intensity. The width of the gaussian is determined by the point-spread function of the imaging system, which the software estimates from the objective numerical aperture and the camera pixel size.
3. Once identified, all diffraction-limited objects in the master channel are tracked by the software to determine when they appear and disappear from the field of view during imaging. From these data, the “lifetime” of each diffraction limited structure in the master channel is calculated.
4. For each of these trajectories, the intensity of the fluorescence signal in each of the subordinate channels (receptor, particle) are also recorded by fitting a 2-dimensional gaussian function to punctate intensity within a narrow search radius around the punctum in the master channel. When there is significant intensity in the particle channel, we classify the trajectory as being particle-associated.
5. The result of this analysis is a distribution of endocytic event lifetimes at the plasma membrane. By applying a threshold to the signal in the particle channel, a separate distribution can be calculated for those endocytic events that are associated with a vesicle and those that did not. These two distributions can be compared to determine the extent to which particles impacted the dynamics of endocytosis. Importantly, because this analysis ignores all particles that did not associate with an endocytic structure, it provides limited insight into the fates of particles with different properties. Therefore, in the next section, we will describe how to modify the analysis to gain more insight into particle fate.

### Analyzing Particle Trajectories to Differentiate Among Possible Outcomes

To analyze particle fate, we need to run a similar analysis to the one described above, but now we will track the particles rather than the endocytic machinery. Similar to how the image sequences were loaded into CMEanalysis to detect puncta in all channels, we will reanalyze these same image sequences here. This step is similar to the *Tracking Endocytic Dynamics During Particle Uptake* section, but the particle channel should be the “master” channel now, with all other channels being the “subordinate” channels. There are four possible outcomes for tracked particles that penetrate beneath cells: i) internalization by the clathrin-coated structure, ii) dissociation of the particle from the clathrin-coated structure without the particle becoming internalized, iii) never associating with a clathrin-coated structure, and iv) internalization by alternative, unmarked endocytosis pathways. Here we describe how to design data filters that isolate with reasonably high confidence particles that were internalized through the clathrin pathway.

1. The first filter that we apply stipulates that a punctum in the particle channel (master) and a punctum in the endocytic channel (subordinate) must be colocalized across a significant number of frames. This criterion eliminates random associations, isolating true colocalization. For this purpose, we use a function that is built into CME analysis, which calculates the statistical confidence that two channels are colocalized. Tracks with greater than 95% confidence of colocalization between the particle (master) and endocytic marker (subordinate) channels are designated as tracking particles that are associated with clathrin-mediated endocytic structures.
2. The next criterion identifies endocytic structures that were present for a sufficient period of time to successfully capture and internalize a particle. We empirically found that these endocytic structures should contain a minimum of 3 frames of at least 60% of the maximum endocytic structure intensity recorded throughout the track. If a frame rate of 2-3 seconds is used, this criterion requires an endocytic structure to colocalize with the particle for at least 6-9 seconds. This criterion is intended to eliminate tracks in which the association between particles and endocytic structures is too brief to result in internalization of the particle.
3. The next criterion is aimed at determining whether particles and endocytic signals disappear at nearly the same time, as would be expected during particle internalization by an endocytic structure. This criterion stipulates that the endocytic structure channel possesses a signal decay of at least 50% of the maximum endocytic structure intensity within at least two frames of a corresponding decline in the intensity of the particle channel. This two-frame threshold should allow for a buffer between four and six seconds (for a two and three second frame rate, respectively). This buffer is needed owing to the intracellular endocytic structure signal which may disappear from the evanescent TIRF field prior to the particle signal. Additionally, this buffer accounts for differences in the sensitivity of the TIRF system to the two fluorescence channels, which can impact the timing of signal loss in each channel.
4. The fourth criterion mandates that the only drop in fluorescence intensity in the endocytic channel is that outlined in step 5. No additional drops in intensity were allowed beyond the drop in step 5. To apply this criterion, we concatenated five additional buffer frames to the end of each track. The absence of additional rises and falls of fluorescence intensity in these frames ensures that the particle has truly been internalized, rather than dissociating from the endocytic site. As such, this criterion increases confidence that we are studying a single particle internalization event by a single endocytic structure.
5. The last filter requires that all endocytic structure signal in frames after the intensity drop are not above 25% the maximum intensity of the endocytic structure channel. This filter eliminates erroneous classification of internalized objects that:
6. may be too close to the noise threshold, or (ii) associated with an endocytic site during a period when its signal fluctuated rather than being internalized by the structure.

## EXPECTED OUTCOMES

Using the above protocol, we sought to determine the impact of particle stiffness on the probability and efficiency of endocytic uptake through the clathrin pathway. To attract particles to the plasma membrane of cells, we engineered a synthetic targeting system that leveraged biotin-streptavidin interactions. Briefly, we used a transferrin receptor chimera, that possesses the cytosolic and transmembrane domains of transferrin, a transmembrane receptor that is constitutively internalized by the clathrin-pathway (Liu et al., 2010). On the ectodomain of our transferrin receptor chimera, we included mEGFP as a fluorescent marker, and a stable monomeric-streptavidin domain for interaction with biotin-conjugated particles (Lim et al., 2011). This model receptor has been described previously in our published work (Ashby et al., 2023). Through the inclusion of biotinylated lipids in liposome compositions, and biotin functionalized polystyrene beads, we targeted delivery of model particles to the plasma membrane of cells which transiently expressed the model receptor. Therefore, our first step was to validate the expression of the model receptor at the plasma membrane.

Successful trafficking of the chimeric receptor to the plasma membrane can be verified using TIRF microscopy (Figure 4B), where the receptor will appear in puncta that colocalize with puncta in the endocytic marker channel. Observing these features confirms: i) successful receptor expression, ii) successful specimen preparation, and iii) proper TIRF imaging conditions, (Figure 4C-E). Once particles are applied, they should be able to penetrate beneath adherent cells within about fifteen minutes. TIRF imaging can then be used to assess the density of particles beneath the cells (Figure 4C-E) (*Cell and Vesicle Imaging* section). A best effort should be made to use groups of cells with similar levels of receptor expression for all experiments that will be directly compared. Expression level can be estimated by quantifying the intensity of puncta in the receptor channel (A values) and the intensity of the plasma membrane surrounding these puncta (C values), (Figure 4F). Among cells with similar expression levels, the density of particles beneath cells can be compared across different particle types. In Figure 4G, we show that liposomes consisting of DOPC, which are less rigid, and DPPC, which are more rigid (Et-Thakafy et al., 2017), penetrate similarly beneath cells, whereas biotinylated polystyrene beads (most rigid) penetrate more poorly, likely owing to aggregation of poly(styrene) particles in the media, as observed at the edges of the cells.

**Figure 4.**
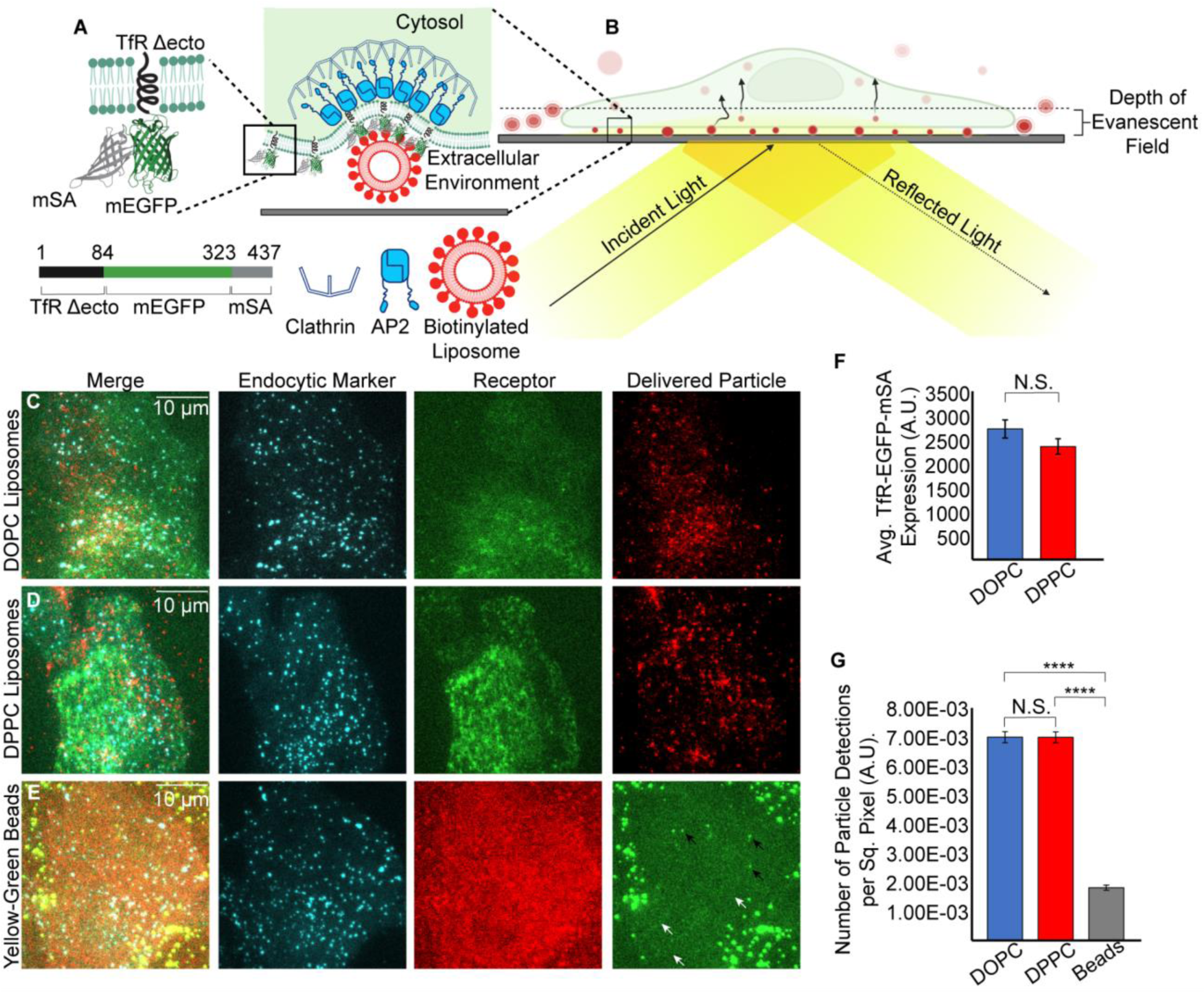
Engineered target receptors facilitate drug-carrier penetration beneath the basolateral membrane. **A)** An engineered model receptor containing a transferrin cytosolic and transmembrane domain conjugated to a monomeric EGFP and monomeric streptavidin to enable targeting of biotinylated lipids. (Created with BioRender) **B)** Schematic of imaging of delivered model drug carriers using TIRF microscopy. **C-E)** Representative images of multiple particle populations delivered to SUM159 cells containing AP2 (cyan) (C-E), a TfR-mEGFP-mSA (C-D), delivered liposomes containing DOPC or DPPC, C and D respectively, Yellow-Green biotinylated beads (E), and a TfR-mRFP-mSA engineered target receptor (E). The black arrows point to aggregated bead complexes and the white arrows point towards small individual beads. **F)** Membrane expression level of the TfR-mEGFP-mSA target receptor in the presence of DOPC and DPPC liposomes. Significance between groups was determined using a student’s t-test (DOPC vs DPPC-p = 0.150). The receptor expression for bead delivery was omitted here owing to the different fluorophore utilized where direct comparisons could not be made. **G)** Normalized number of particle detections beneath cells for the DOPC liposome, DPPC, liposome and Biotinylated Yellow-Green bead populations. Significance between groups was determined using a student’s t-test (DOPC vs DPPC-p = 0.976; DOPC vs Bead-p < 10-5; DPPC vs Bead-p < 10-5) Data for F and G were acquired over at least 3 independent trials with at least 12 cells analyzed per trial culminating with N= 84, 109, and 110 cells for the DOPC, DPPC, and Yellow-Bead delivery conditions respectively.

TIRF images of the cells also reveal colocalization events between particles and clathrin-coated structures. Figure 5A shows a cell (white dashed line) to which liposomes consisting of DOPC, labeled by Texas Red, were delivered. The inset of Figure 5A shows a snapshot of a colocalization event between a clathrin-coated structure (cyan) and a liposome (red).

**Figure 5.**
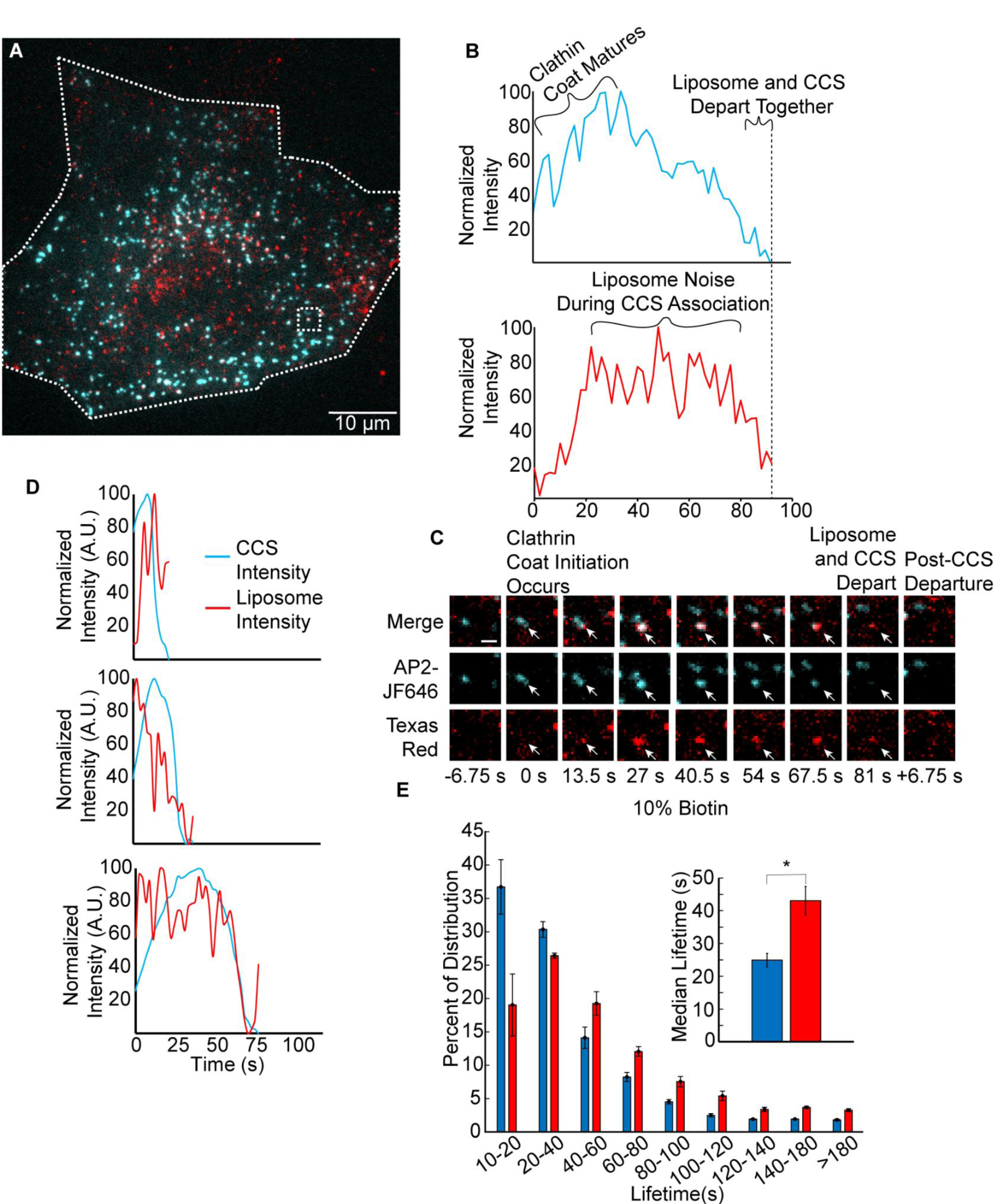
Tracking the endocytic machinery enables the observation of overall particle effects on the dynamics of endocytosis. **A)** Representative image of a SUM159 cell exposed to DOPC liposomes containing 10% biotinylated lipids. Adaptor protein 2 (cyan) is represented at the cytosolic leaflet of the plasma membrane, and liposomes are free to diffuse between the basolateral membrane and the coverslip. Colocalization (appears as white) between the 2 channels over statistically significant time scales denotes colocalization that can be analyzed. The periphery of the cell is highlighted with the dashed white line, and the dashed box represents a representative colocalization event. **B)** Representative intensity profile of the growing clathrin-coat and its colocalization with the liposome that is shown in the dashed white box in A. **C)** Montage of the growing endocytic site shown in the dashed box in A and the intensity profile in B that show the simultaneous departure of a liposome and clathrin-coat indicating successful internalization. The scale bar is 1 µm. **D)** Representative time cohorts for the DOPC liposome delivery group that show over long-time scales there is good agreement for the simultaneous intensity decrease of the liposome intensities and the clathrin-coats averaged into cohorts of 10-20s, 20-40s, and 60-80s. Average cohorts were composed of 981, 948, and 408 individual events for 10-20s, 20-40s, and 60-80s cohorts respectively. **E)** Representative time cohorts for clathrin-sites that associate with a liposome (red) and those that do not (blue). The median lifetime inset indicates a longer lifetime for the clathrin-sites that interact with a liposome in comparison to those that do not. Error bars represent the standard error of the mean across three independent trials. Data for E was acquired over at least 3 independent trials with at least 12 cells analyzed per trial culminating with N= 84, cells analyzed that were exposed to DOPC containing liposomes.

Using the protocol describe above, these events can be tracked individually through time as shown in Figure 5B, where the liposome and clathrin-coated structure disappear simultaneously from the TIRF field, suggesting internalization of the particle by endocytosis (*Analyzing Particle Trajectories to Differentiate Among Possible Outcomes* section). Figure 5C shows a montage of the intensity trace in Figure 5B and highlights this phenomenon.

As outlined in the *Tracking Endocytic Dynamics During Particle Uptake* section, CMEanalysis was used to track all endocytic events. We first tracked the endocytic marker channel (AP2-JF646, cyan) and filtered for events that displayed colocalization with the subordinate liposome channel (red). From here, we averaged the intensities of events that occurred over similar timescales to create plots which show the average intensity trajectories over time, Figure 5D. These trajectories show the rise, plateau, and subsequent fall in fluorescence intensity in both the endocytic marker and particle channels. The literature on the dynamics of clathrin-mediated endocytosis collectively suggests that short-lived events (<20 seconds) are often abortive, disassembling stochastically rather than leading to productive endocytosis (Ehrlich et al., 2004; Loerke et al., 2009). This is likely why the intensity in the liposome channel does not decrease simultaneously with the intensity in the AP2 channel at the end of the trajectories that make up the shortest time cohort.

However, when we study longer-lived events (20-40 seconds, and 60-80 seconds), we see simultaneous intensity declines, suggesting that many of these events result in particle internalization. When we split all endocytic events into those that are associated with a particle and those that do not, we can see that particles tend to associate with longer-lived endocytic structures (Figure 5E). Our prior work (Ashby et al., 2023) suggested that this effect arises from longer-lived structures having a higher probability of colliding with a particle during their limited lifetime at the cell surface.

## QUANTIFICATION AND STATISTICAL ANALYSIS

In the *Expected Outcomes* section, we explain how the effects of particle delivery on endocytosis and vice versa can be analyzed through the tracking endocytic structures and filtering based upon the presence or absence of a delivered particle. As shown in the *Analyzing Particle Trajectories to Differentiate Among Possible Outcomes* section, we can also employ this analysis technique on the particles themselves. When we follow the steps highlighted in the *Vesicle Imaging, Vesicle Detection,* and *Vesicle Intensity to Diameter Conversion* sections, we create a conversion factor for particle intensity to particle diameter that can be applied to particles that were tracked according to the *Analyzing Particle Trajectories to Differentiate Among Possible Outcomes* section. This analysis reveals that the diameter distribution for liposomes consisting of DOPC and DPPC that were able to penetrate beneath cells was not very different from the initial diameter distribution. In contrast, for polystyrene beads, which are more rigid than liposomes, there was a significant shift toward larger diameters, likely because beads tended to cluster together in the cell culture environment.

When the filters described in *Analyzing Particle Trajectories to Differentiate Among Possible Outcomes* are applied, we can identify groups of particles that penetrated beneath cells, associated with endocytic structures, and were internalized by endocytic structures, where each subsequent group is a subset of the former group. Figure 7A-B shows the results of this analysis by plotting the distribution of particle diameters for: particles prior to introduction to cells (A), particles that penetrated beneath cells (B, repeated from Figure 6A-C), particles that associated significantly with endocytic structures (C), and particles that were internalized by endocytic structures (D). Notably, there is a shift toward smaller diameters for particles that associate with endocytic structures (Figure 7C) and again for particles that are internalized (Figure 7D). These results suggest that the internalization process favors small particles, perhaps because they represent a smaller mechanical barrier to endocytosis. This trend becomes even clearer when we plot the probability of association with an endocytic structure for particles with diameters less than 40 nm, relative to the overall probability for all particles in the distribution, Figure 7E. Similarly, Figure 7F plots the same relative probabilities for the internalization step, comparing those particles with diameters below 40 nm to the full population. These data show that the probability of association and internalization are both substantially increased for the smallest particles in the distribution. Interestingly, the fractional increase is smallest for the softest particles (DOPC liposomes) and largest for the hardest particles (polystyrene beads), with the particles of intermediate stiffness (DPPC liposomes) displaying an intermediate effect. These trends suggest that as particle stiffness increases, endocytic uptake requires that particles be smaller and smaller to be efficiently internalized.

**Figure 6.**
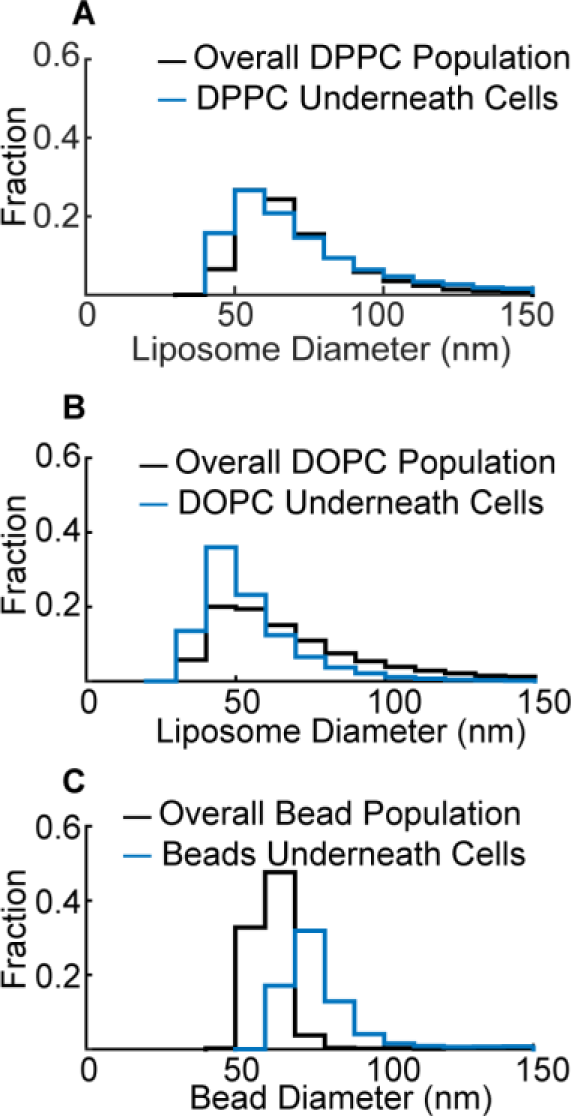
Comparisons for various particle types enable judgment on how physical particle properties can dictate particle-cell interactions. **A-C)** Overall particle populations (black) and the distribution of particles present beneath cells for DPPC liposomes (A), DOPC liposomes (B), and biotinylated beads (C). The black curves are created from converted tethered vesicle distributions (as seen in Figure 2) and contain 8,960, 47,502, and 6,245 particle puncta for DPPC, DOPC, and beads respectively. The blue curves are taken beneath cells across 3 independent trials with a minimum of 12 cells analyzed per trial for a total cell count of 102, 84, and 110 cells for DPPC, DOPC, and beads respectively. The particle totals for each condition are 17,502 DPPC liposomes, 26,537 DOPC liposomes, and 6,094 beads.

**Figure 7.**
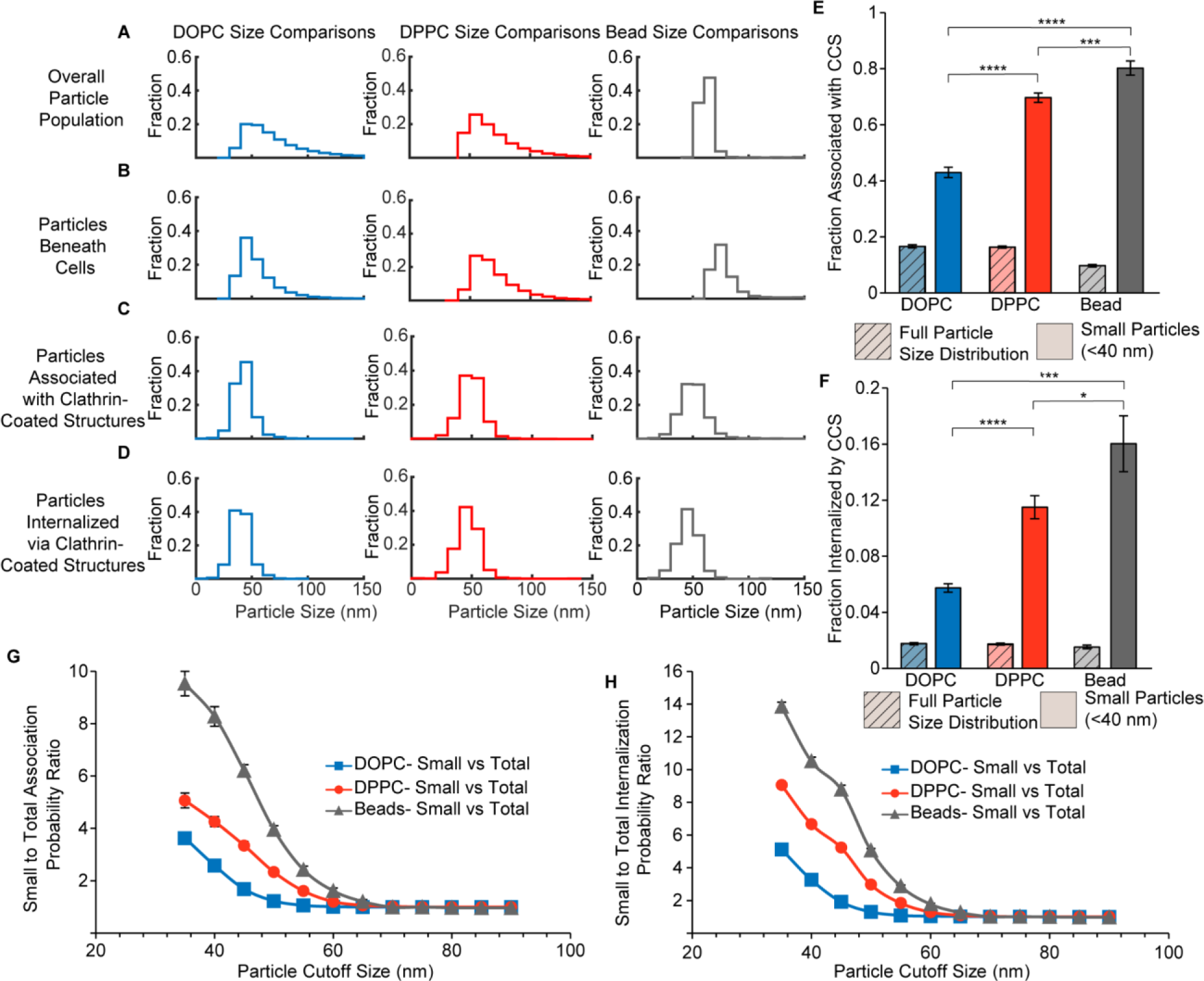
Determining successfully internalized particles and elucidating the effects of particle stiffness. **A-D)** Size distributions of particle diameters containing either DOPC (blue), DPPC (red), or biotinylated beads (gray). The top plot is the overall size distributions of the particles prior to delivery to cells (particle counts were 47,502, 13,318, and 6,245 for DOPC, DPPC, and beads respectively). The second plot is the distribution of particle diameters that penetrated beneath the basolateral membrane of adherent cells (particle counts were 26,537, 21,788, and 6,094 for DOPC, DPPC, and beads respectively). The third plot is the distribution of particle diameters that penetrated beneath adherent cells and associated with a clathrin coated structure (particle counts were 15,152, 14,730, and 2,028 for DOPC, DPPC, and beads respectively). The fourth plot is the distribution of diameters for particles that penetrated beneath adherent cells, associated with a clathrin-coated structure, and were successfully internalized by that structure (particle counts were 1,797, 1,540, and 358 for DOPC, DPPC, and beads respectively). **E)** Bar graph representing the probability of any particle associating with a clathrin coated structure (hashed group) compared to those that were beneath a 40 nm diameter cutoff (solid) for DOPC containing liposomes (red), DPPC containing liposomes (blue), and biotinylated beads (gray). The significance of differences between the small particle groups was determined by using a student’s t-test assuming unequal variances (DOPC vs DOPC-p < 10-5; DOPC vs Bead-p < 10-5; DPPC vs Bead-p = 0.0008) **F)** Bar graph representing the probability of any particle becoming successfully internalized by a clathrin-coated structure (hashed group) compared to those that were beneath a 40 nm diameter size cutoff (solid) for DOPC containing liposomes (red), DPPC containing liposomes (blue), and biotinylated beads (gray). The significance of differences between the small particle groups was determined by using a student’s t-test assuming unequal variances (DOPC vs DOPC-p < 10-5; DOPC vs Bead-p < 10-5; DPPC vs Bead-p = 0.0410) For E and F, at least 3 independent trials were conducted for each condition with a minimum of 12 cells analyzed per trial. The total number of cells analyzed was 84, 109, and 110 for DOPC, DPPC, and biotinylated beads respectively. **G-H)** The ratio of the probability of a particle beneath a diameter-cutoff (x-axis) of associating with a clathrin-coated structure (G), compared to the probability of the full particle diameter size distribution, and the ratio of the probability of a particle beneath a diameter size-cutoff of becoming successfully internalized by a clathrin-coated structure, compared to the probability of the full particle diameter distribution. The total number of cells in G and H were 68, 92, and 44, for DOPC (blue), DPPC (red), and biotinylated beads (gray) respectively. Any cells containing fewer than 150 internalization events beneath a 35 nm diameter cutoff were excluded from this analysis.

To further probe this phenomenon, we varied the particle diameter cutoff (x-axis) to assess the probability of association with endocytic structures and internalization for particles beneath the cutoff in comparison to the probability for the entire distribution of particles, Figure 7G, H. These plots extend the analysis in Figures 7E, F to all possible diameter thresholds, where the value on the vertical axis indicates the relative preference of endocytic structures for particles with diameters below the threshold on the horizontal axis. Here we see that, for the stiffest group of particles (polystyrene beads), the smallest particles in the distribution are more than 10-fold more likely to be internalized than the average particle in the distribution. Similar trends exist for particles of intermediate stiffness (DPPC liposomes) and lowest stiffness (DOPC liposomes), where the magnitude of the trend decreases with decreasing particle stiffness. Overall, these data demonstrate that particle size and stiffness are convoluted during endocytic uptake. While endocytic structures prefer particles of small diameter, regardless of their stiffness, the requirement for small size becomes increasingly severe as particle stiffness increases. We speculate that this trend arises from the ability of softer particles to deform during their interactions with cells, making it easier for them to diffuse to endocytic sites and fit within growing endocytic structures, such that they become internalized. On the basis of these data, researchers who design particles for endocytic uptake should keep in mind that rigid particles need to be very small in diameter to achieve efficient uptake. Importantly, these insights have arisen from examining heterogeneity in diameter across particle populations, an analysis which is not possible using traditional batch techniques to examine particle uptake by endocytosis.

## ADVANTAGES

Some advantages of this method include the opportunity to quantify particle internalization by a specific endocytic pathway. In contrast, batch techniques, like flow cytometry and Western blot analysis, quantify total uptake by all endocytic pathways. While pharmacological inhibitors can be used to narrow the range of pathways participating in internalization, their impact is imprecise, and they typically have off-target effects (Basagiannis et al., 2021; Park et al., 2013). Additionally, through the tethered vesicle assay and associated diameter conversion, we can estimate the diameters of individual particles that are being internalized. This information allows the user to determine how the efficiency and dynamics of internalization vary with particle diameter within a heterogeneous distribution of particles. Using this approach, hundreds of particles, each with their own diameter, can be tracked in parallel, providing statistics on the probability of uptake for particles with different properties. Owing to the inherent heterogeneity of many particle manufacturing protocols (Gkionis et al., 2020; Larsen et al., 2011), this information can provide unique insights into uptake of particles by endocytosis. In contrast, bulk methods report the overall uptake of heterogeneous particles, providing no direct insight into the impact of heterogeneity on uptake. Lastly, TIRF microscopy restricts analysis to particles that penetrate beneath cells. This requirement selects for particles that have the potential to penetrate into tight spaces between cells in tissue, more accurately recapitulating the in vivo setting and potentially foreshadowing delivery challenges that are not well captured in bulk assays, where most particles likely enter cells from their more accessible apical surface.

## LIMITATIONS

The restriction of this technique to uptake events that occur on the basolateral membrane may be regarded as a limitation by researchers who are interested in uptake by cells in which the apical surface plays an important role in delivery, such as endothelial cells, and cells that mimic epithelial barrier layers such as the gut and lung epithelia. Additionally, while this method can track hundreds of internalization events per cell and thousands per imaging session, the throughput remains relatively low compared to batch methods. For example, flow cytometry can be performed on tens of thousands of cells per measurement. Our method also requires that particles be densely labeled with relatively bright, stable fluorophores that resist photobleaching, such that the particles can be tracked over several minutes of continuous imaging.

## OPTIMIZATION AND TROUBLESHOOTING

### Improper Sample Slide Preparation

It is critical that the imaging slide be prepared in such a way that the coverslip is almost perfectly parallel to the slide. In this way, when the slide is held by the microscope stage, the coverslip will be held perpendicular to the objective, a key requirement for TIRF microscopy. When this requirement is not met, it is often impossible to focus on the cellular plasma membrane during TIRF illumination. Common causes of improper sample slide preparation include a wrinkle in the layer of double-sided tape that is used to adhere the coverslip to the slide and/or improper application of valap, such that the valap layer contacts the objective. Stretching the tape while sticking it to the coverslip and applying a conservative amount of valap can help to address this issue.

### Improper Cell Heating

Improper heating of the cells during imaging can lead to a misrepresented distribution of endocytic lifetimes, as the assembly of endocytic proteins is a function of temperature. This problem is indicated by an endocytic structure lifetime distribution that show shifts towards higher fractions of either abortive (too warm) or stalled events (too cold) than would be present during observation of the clathrin-mediated endocytosis pathway at physiological temperature. These issues can be eliminated by determining the time required for your sample holder to reach a steady temperature and waiting to begin the imaging session until that time has elapsed.

### Particles Do Not Penetrate Beneath the Cell

Another potential failure mode that may occur when following this method is that particles may not penetrate beneath adherent cells. If so, the absence of particles will be obvious once imaging commences. Cells will appear outlined by the particles, which may stick to the glass coverslip that the cell is growing on. When too few particles penetrate beneath cells, there will likely be too little data to extract meaningful trends and conclusions. Failure of the particles to penetrate beneath cells typically occurs for two reasons: (i) the particles are not small enough to penetrate between the adherent cell membrane and the coverslip, or (ii) there is a component in the particles that is interacting with the glass coverslip, causing the particles to adhere to the glass, rather than penetrating beneath cells. Particles with diameters larger than 100 nm are likely too large to penetrate efficiently beneath cells. Dynamic light scattering can be used to assess the particle size distribution, helping to determine what fraction of the particles are likely to penetrate. Sonication of particles can help to break up clusters that would be too large to penetrate. If particles adhere to the coverslip, we recommend coating them with hydrophilic molecules such as polyethylene glycol chains. Notably, plasmid DNA used to transfect cells with fluorescent markers can adhere to the coverslip, where it may attract particles, owing to its strong negative charge. Washing the cells thoroughly after transfection can help to minimize this effect.

## SAFETY CONSIDERATIONS AND STANDARDS

While executing this method, standard, biosafety level-2 (BSL-2) laboratory procedures should be used, as is the case for all experiments with mammalian cells. During particle preparation, organic solvents such as chloroform used to dissolve lipids should be handled in a chemical fume hood and disposed of in a safe manner according to institutional and governmental regulations. If liposomes are being created as a model drug-carrier and they will be broken into smaller diameter sizes via probe sonication, hearing protection is recommended during sonication.

## CONNECTIONS

Endocytosis is a classic example of membrane curvature induction in cells, and many researchers study the energetics of endocytosis and how this may affect membrane structure in theoretical, computational, and experimental models. While this method provides a guide towards studying particle uptake, studying the energetics of particle uptake as a steric barrier to membrane bending, and how a curved substrate affects membrane coupling and organization are potential expansions of this work. In this double volume, there are many methods that may provide further insight towards the forces at play during particle internalization that could facilitate further expansion of this work.

These other potential methods include a computational model by Peter Tielman’s group titled, “Analyzing curvature and lipid distributions in molecular dynamics simulations of complex membranes”, a computational model by Reinhard Lipowsky’s group titled, “Multiscale remodeling of biomembranes and vesicles”, and an experimental model from Nick Brooks group titled, “Effects of lateral and hydrostatic pressure on membrane structure and properties.”

